# Ethnobotanical and nutritional study of *quelites* sold in two traditional markets of Oaxaca, Mexico

**DOI:** 10.1101/453225

**Authors:** G.I. Manzanero-Medina, A. Pérez-Herrera, H. Lustre-Sánchez, M.A. Vásquez-Dávila, N.F. Santos-Sánchez, M.A. Sánchez-Medina

## Abstract

**Background.:** In Mexico, it is called *quelites* to certain edible vegetables (young plants, germ, shoots or flowers). Since pre-Hispanic times, *quelites* have been eaten as a source of vitamins, minerals and proteins. Now, its traditional and healthy consumption has decreased. We studied the *quelites* of two traditional markets in the Valles Centrales of Oaxaca state, Mexico using an ethnobotanical and nutritional approach.

**Methods:** From July 2017 to July 2018, weekly ethnobotanical interviews were conducted with 26 collectors-sellers of the Zimatlán market and 36 in the Zaachila market. The vegetal supply was acquired, herborized and identified by through dichotomous keys. There were determined the proximal composition, phenolic compounds, flavonoids, antioxidant capacity and mineral content of the floral structures of two *quelites’* types. The statistical analysis was performed through a one-way analysis of variance (ANOVA) of Tukey HSD.

**Results:** In two sampled markets, 23 species belonging to 11 botanical families were registered, from which leaves, branches, stems, flowers and fruits are eaten. The flowers of the species *Diphysa americana* (Q1) and *Phaseolus coccineus* (Q2) are the most used for human consumption of the communities involved in the sale of the sampled *quelites*. Both flowers had important amounts of proteins (2.66-3.29%) and fiber (1.66-2.43%). Q1 had higher content of phenols and flavonoids and therefore higher antioxidant capacity than Q2 (p <0.05). When we talk about Q2 minerals, it presented a greater amount of Zn, Ca and Mg in comparison to Q1 (p> 0.05).

**Conclusions:** In local markets of the state of Oaxaca, a wide variety of *quelites* are usually found, where their botanical structures, such as flowers, are widely eaten. The flowers of Q1 and Q2 proved to be a rich source of proteins and bioactive compounds, as well as minerals. Showing thus to be a food alternative to enrich the human diet.

## INTRODUCTION

Located in southern Mexico, the state of Oaxaca has a high diversity of vegetation types and a great floristic richness of more than 30,000 [1–3]. In this range of biodiversity there are sixteen ethnic groups with extensive traditional knowledge on the use and management of biotic resources. Due to its extension and complexity, it still does not have a finished and updated potential flora [4]. In Mexico, the knowledge of native flora by ethnic groups allowed, since pre-Hispanic times, the use of diverse vegetal resources. This is the case of *quilitl*, a word of *Náhuatl* origin which is translated in Spanish as *quelites* and it is edible tender vegetables, young plants, germ or shoots of some trees and, in some cases, edible flowers [5].

Currently, the diet pattern of a large part of the mexican population is featured by industrialized food products and the gradual reduction of traditional foods. This transition in eating behavior, with a clear trend towards the consumption of industrialized products with a high energy content, has given rise to the appearance of chronic noncommunicable diseases such as cardiovascular diseases and type 2 diabetes mellitus. However, at the same time it begins to recognition and return to the consumption of non-industrialized foods, mostly with local and / or regional identity [6–8].

In Mexico, the vegetal diversity with the potential for use includes around 5000 species from different botanical families [9], most of them are herbaceous and wild [10], with food and medicinal uses [11], mainly sold in local and regional markets of the south-southeast region. The use of a large number of vegetal species has a pre-Hispanic origin and its consumption still continues in rural communities where there is a wide variety of foods for local cuisine, which has been little studied. Within these foods are the *quelites*, from which can be used the leaves, the flower and/or the fruit. In Mexico are consumed around 500 species chosen by the local traditions of the different peoples and regions [12]. *Diphysa americana* (Mill) M. Sousa and *Phaseolus coccineus* (L.) are two types of *quelites* where their floral structures are consumed in the rural communities of the Valles Centrales’ region of Oaxaca. *Diphysa americana* known as *flor de gallito*, *guachipelín*, *yaga-yetzi* (in Zapotec language), among other names, is a vegetal species that extends from Mexico to Panama and is a slow growing deciduous tree that reaches from 20 to 65 ft tall [13]. *Phaseolus coccineus*, known as *quelite*, *frijol de monte*, ayocote, colorín, among other names, is a perennial and climbing plant of several feet long which is spread along Mesoamerica and it’s a legume native of the area of the current countries of Mexico and Guatemala [14].

For centuries, edible flowers have been an integral part of human nutrition. For example, in central Europe the black elder flowers (*Sambucus nigra*) fried or battered are common, and the dandelion flowers boiled with sugar were used as *ersatz honey*. In addition, the flowers were used to decorate foods prepared for the feasts and banquets of the noble people [15]. In China and Japan, edible flowers have been eaten for thousands of years [16, 17].

Because *quelites* and their botanical structures are considered under-used plants in Mexico, it is necessary to resume their study to incorporate them into the Mexican’s current feeding pattern. For this, when talking about balanced diets, a wide variety of foods is included, involving green leaves and flower structures. The content of common components such as proteins, fats and vitamins in flowers are not very different in relation to other organs of the plant such as leaves [18, 19]. It has been found that some flowers are rich in phenolic compounds and have high antioxidant activity, as well as the ability to inhibit chronic diseases [20–22]. In addition, the color they got is predominantly due to several chemical compounds such as carotenoids and flavonoids [23]. The latter ones are correlated to their high antioxidant capacity [24]. In this context, the purpose of this paper was to document ethnobotanical knowledge about *quelites* and nutritional research of two best-selling species in two traditional markets of Zaachila and Zimatlán in the region of the Valles Centrales of Oaxaca. The market of Zaachila is one of the most diverse of those located in the Valles Centrales’ region; as well as that of Zimatlán. Considering the aforementioned, these two markets were chosen in *Plaza* Days since they are a focal point for trade within the Valley of Oaxaca.

## METHODS

### Sampling area

The state of Oaxaca is located at southern Mexico and is divided into 8 socio-cultural regions. The study markets are located in the region of the Valles Centrales where the city of Oaxaca de Juárez is located and has 121 municipalities. In the municipality of Zimatlán de Álvarez, on Wednesdays, the local market is established on the streets, while on Thursdays it is sitted in the Villa de Zaachila municipality, **Figure 1**.

**Figure 1.**
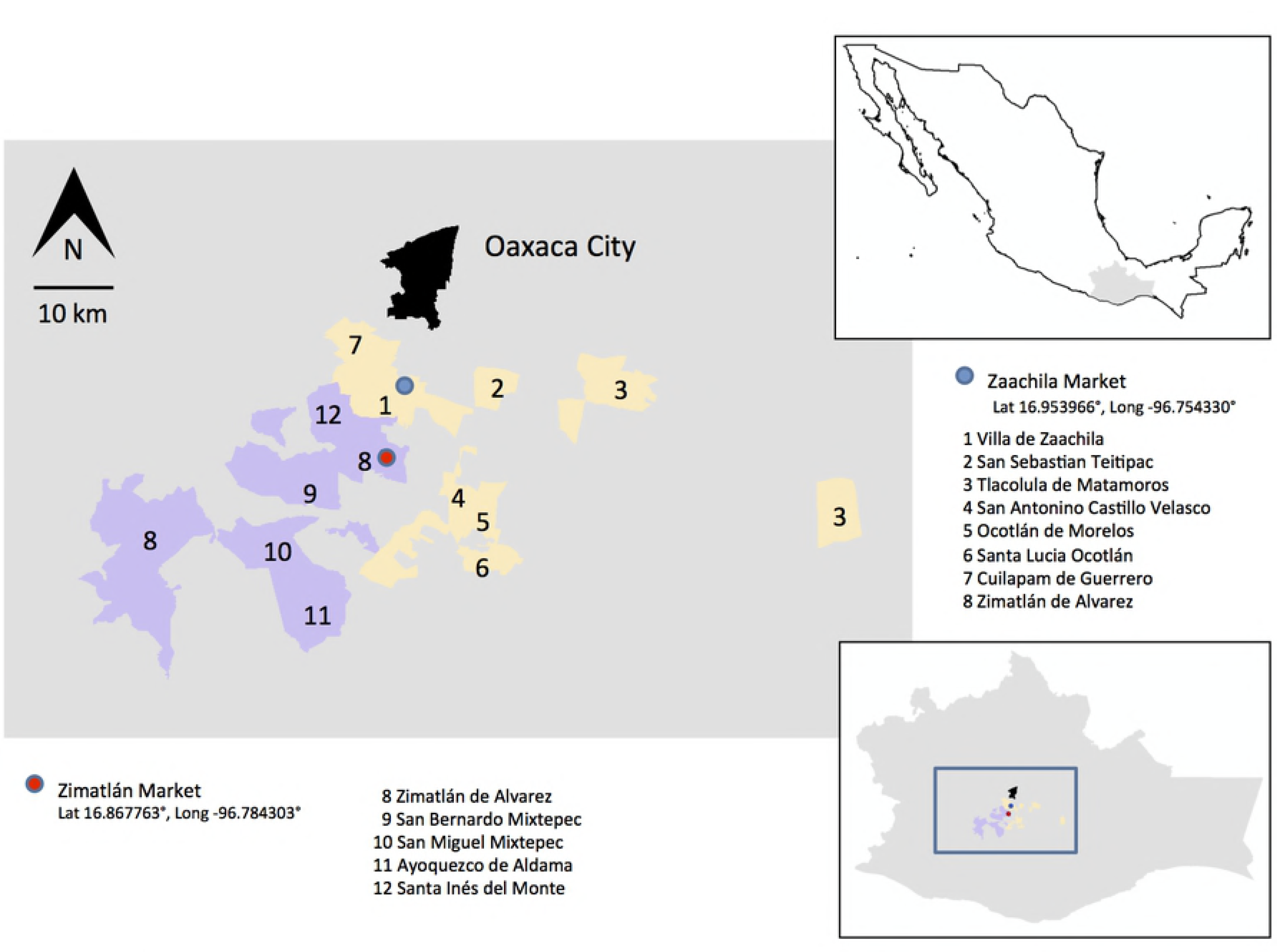
Location map of the two sampled markets and sites involved in the sale of *quelites* from the Valles Centrales’ region, Oaxaca, Mexico. Map made by Matthias Rös.

The municipality of Villa de Zaachila, is located between an altitude of 1400 and 2300 m, between the parallels 16º52′ and 17º02′ of north latitude; the meridians 96º39 ‘and 96º52′ of west longitude. The range of temperature that exists in Zaachila is of 16-22 ºC, a range of precipitation of 600-800 mm and a dry climate, semi-dry-semi-warm (70.99%), semi-warm-sub-humid with rains during summer (26.72%) and mild-sub-humid weather with rain during summer (2.29%) [25]. The municipality of Zimatlán de Álvarez, is located between an altitude of 1400-3100 m, between the parallels 16º34 ‘and 16º57′ of north latitude; meridians 96º45 ‘and 97º13′ west longitude. The temperature range that exists in Zimatlán is 12-22 ºC, with a range of precipitation of 600-2500 mm and a mild-sub-humid weather with rains during summer, more humid (37.45%), semi-warm-sub-humid with summer rains, more humid (21.88%), semi-warm-sub-humid with rain during summer, less humid (16.88%), mild-humid with abundant rainfall during summer (15.17%), dry, semi-dry-semi-warm (4.99%) and mild-sub-humid weather with rain during summer, medium moisture (3.63%) [26].

### Sampling in the local markets of Villa de Zaachila and Zimatlán de Álvarez

Every week, on Wednesdays and Thursdays’ morning, the markets of Zimatlán and Zaachila were visited, respectively, with the purpose of identifying all the stands where the *quelites* are sold during the *Días de Plaza* in each one of them. Part of the botanical structures was herborized and preserved properly labeled to be stored in the *Herbario OAX* of the CIIDIR-IPN-Oaxaca, through the technique of Long and Chiang [27] which consists of a collection of the samples, drying, labeling and identification of the species.

In order to know the ethnobotanical aspects of the *quelite* species, data was obtained from interviews with sellers about the places of origin and their habitat features, as well as the methods of collection, uses, ways of use and demand of each species [28]. There were taken pictures of the vegetal species with their original presentation in the market. In some cases, the places of origin and collection of the *quelites* were visited, following the invitation and attention of the *quelites’* collectors-sellers.

The information obtained was registered for each sample taken to be incorporated into a database: scientific name, botanical family, indigenous name, name in Spanish, structure used, use, way of use, origin, harvest time, geographical distribution, vegetation or habitat type, origin and degree of management (grown, wild, tolerated) [28]. To perform the chemical and nutritional analysis, previously were selected the floral structures of the *Diphysa americana* and *Phaseolus coccineus* species (**Figure 2**).

**Figure 2.**
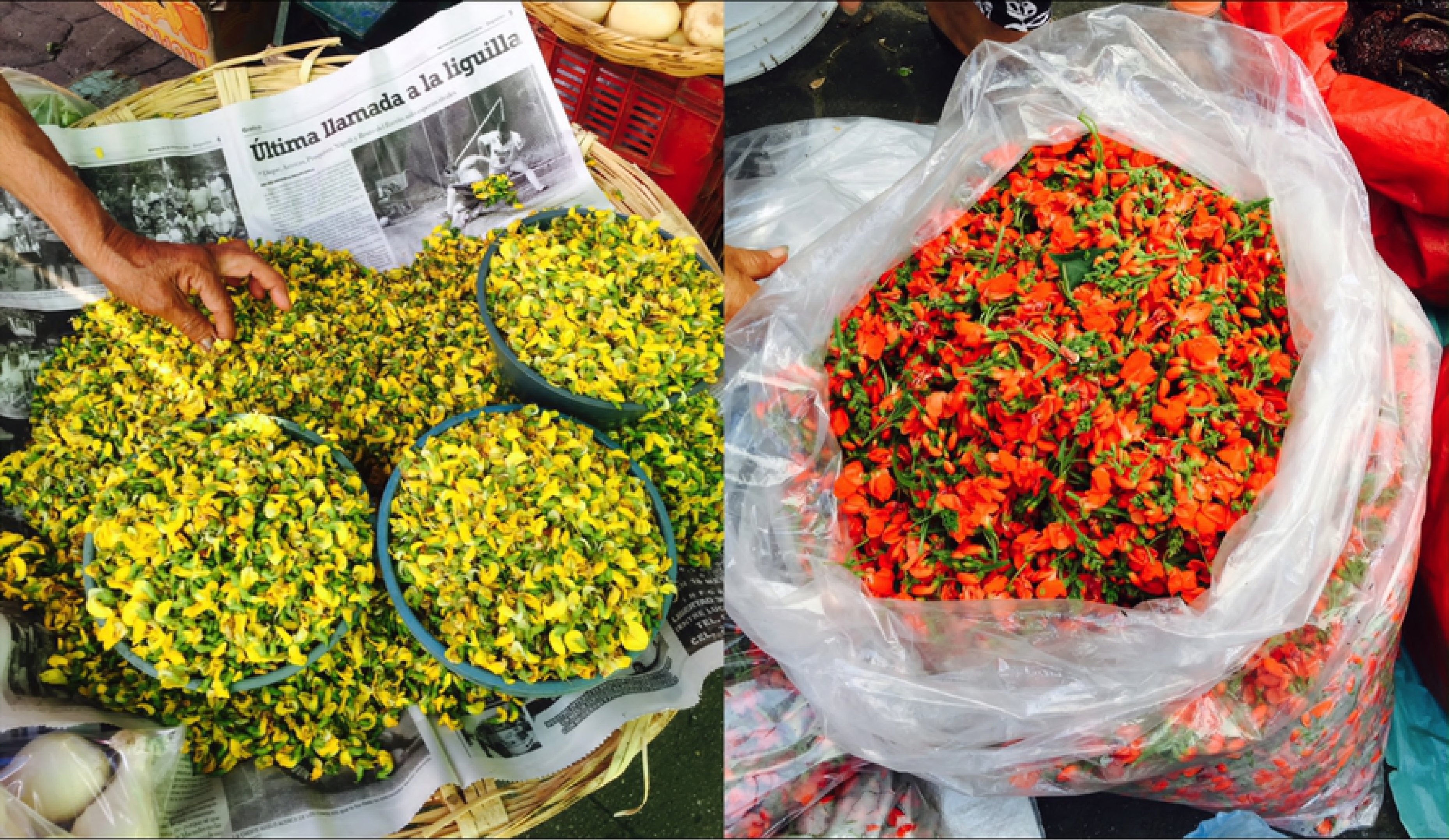
*Diphysa americana* and *Phaseolus coccineus* flowers (from left to right in the picture) collected in the local markets of Villa de Zaachila and Zimatlán de Álvarez.

### Nutritional analysis

#### Moisture, ashes, proteins, fats, fiber, carbohydrates and energy

For moisture, ash, fat, protein and fiber determinations were used AOAC methods 925.09, 923.03, 920.39, 954.01, 962.09, respectively (2000) [29], in triplicate for each sample of *quelites*. The moisture was determined by drying at 105 °C until constant weight. The ashes of the remanent were determined by calcination at 550 °C for 8 h. The nitrogen was obtained by the Kjeldahl technique and the concentration of the protein was calculated using the nitrogen factor of 6.25. The content of fats was determined by extracting them with the petroleum ether, and the total dietary fiber was determined by the incineration by the organic state that remains by the digestion with solutions of sulfuric acid and sodium hydroxide. The carbohydrates, nitrogen-free extract (NFE), were determined by subtraction (100 less the sum of proteins, fats, moisture, ashes and dietary fiber). The total energy was based on the results of carbohydrates, proteins and fats. The conversion factors were based on the metabolizable energy and analytical methods: 16.7 kJ/g for carbohydrates, 16.7 kJ/g for protein and 37.7 kJ/g for fats [30].

#### Preparation of the extract

Extracts of the *Diphysa americana* (Q1) and *Phaseolus coccineus* (Q2) *quelites* were prepared as described by Rababah et al. [31] with some modifications. About 20 mg of dry sample were mixed in 10 mL of 90% methanol. Subsequently, they were sonicated for 30 minutes and then filtered using Whatman filter paper number 3.

#### Quantification of phenolic compounds

The determination of the phenols was made according to the method using the Folin-Ciocalteu reagent [32]. To the extracts of the two samples of *quelites* (1 mL per triplicate) were added a volume of 1 mL of distilled water, 1 mL of 0.5% Na_2_CO_3_ aqueous solution (w/v) and 1.0 mL of Folin-Ciocalteu 2 N reagent diluted 10 times in water. The samples were vortexed protected from light and allowed to stand for 1 hour at 25 °C. The absorbance was measured at 750 nm in a UV-visible spectrophotometer (Shimadzu UV-1800). The results were compared with a calibration curve (r^2^=0.9814) using gallic acid standards (0-250 ppm). The content of total phenols was determined from the equation of the curve and expressed as mg of gallic acid equivalents (GAE)/g dry matter.

#### Quantification of total flavonoids

The content of total flavonoids was made according to the method described by Chen et al. 2014 [33]. Extracts of the two samples of *quelites* (2 mL in triplicate) were added with a volume of 150 μL of 5% NaNO_2_ and allowed to stand for 5 minutes, then 150 μL of 10% AlCl_3_•6H_2_O were added and after 6 minutes of rest was added 1 mL of NaOH at 1 M. The absorbance was measured at 510 nm in a UV-visible spectrophotometer (Shimadzu UV-1800). The results were compared with a calibration curve (r^2^=0.998) using quercetin standards (0-100 ppm). The content of total flavonoids was determined from the equation of the curve and expressed as mg of quercetin equivalents (QE)/g dry matter.

#### Determination of antioxidant capacity

It was used the technique of the 1,1-diphenyl-2-picrilhydrazil radical (DPPH) (Sigma, Aldrich) to determine the antioxidant activity expressed as percent inhibition, following the technique described by Matthaus [34]. 100 μL of each extract were taken at 5 different concentrations (50, 25, 12.5, 5 and 2.5 mg/mL) and 2.9 mL of DPPH solution (50 mg/100 mL) was added to each. All reactions were carried out for 30 minutes at room temperature. The absorbance at 515 nm was measured against the target (without extract) in a UV-visible spectrophotometer (Shimadzu UV-1800). The antioxidant activity is expressed como mol Eq.Trolox/mg dry matter.

#### Determination of minerals

Samples Q1 and Q2 were dried in an oven at 230°F all night long, this sample was mineralized by dry route (AOAC 923.03) at 600 °C for 24 hours. The ashes were dissolved in 10 mL HCl 20% (v/v) heated for 30 min at slow boiling. The clear solution obtained was filled with 25 mL deionized water in a volumetric flask [35]. Each sample was made in triplicate. For the quantification of the minerals, calibration curves were constructed for each mineral (zinc, copper, iron, calcium and magnesium), according to the requirements of the atomic absorption spectrophotometer GBC 932AA, equipped with an air-acetylene flame burner, and empty cathode lamps for each element as source of radiation [36]. From the solutions obtained for the quantification of minerals in the samples, dilutions were made according to the concentrations required by the calibration curves of each mineral and the readings were made in the atomic absorption spectrophotometer. The target of each sample was obtained with the same treatment used for the sample but without the ashes. For the determination of calcium and magnesium both the samples and the standards contained 5000 ppm of lanthanum as an ionization regulator. Each reading was made in triplicate and the quantity was obtained from the equation of the curve of the standards used.

#### Analysis of results

The data obtained were analyzed with the statistical program SPSS version 15.0 (SPSS Inc., Chicago, IL, USA). All determinations were made in triplicate. The results were expressed as the averages  SD. The statistical analysis was performed through a one-way analysis of variance (ANOVA) of Tukey HSD.

## RESULTS

### Frequency of presence of species considered quelites in the markets of Villa de Zaachila and Zimatlán de Álvarez

26 *quelites* were sampled in the Zimatlán market and 36 in the Zaachila market. In the first market, 15 species of *quelites* belonging to 9 botanical families were registered and in the market of Zaachila there were 23 species grouped in 11 botanical families. The *quelites* that are sold in these markets are consumed both fresh and dried and the main edible parts are the leaves, except for the species selected for the chemical-nutrimental study, where the botanical structure that is eaten are the flowers (**Table 1**). In the two markets studied, a total of 23 species were registered, 15 of which occur in both markets (**Figure 3**).

**Figure 3.**
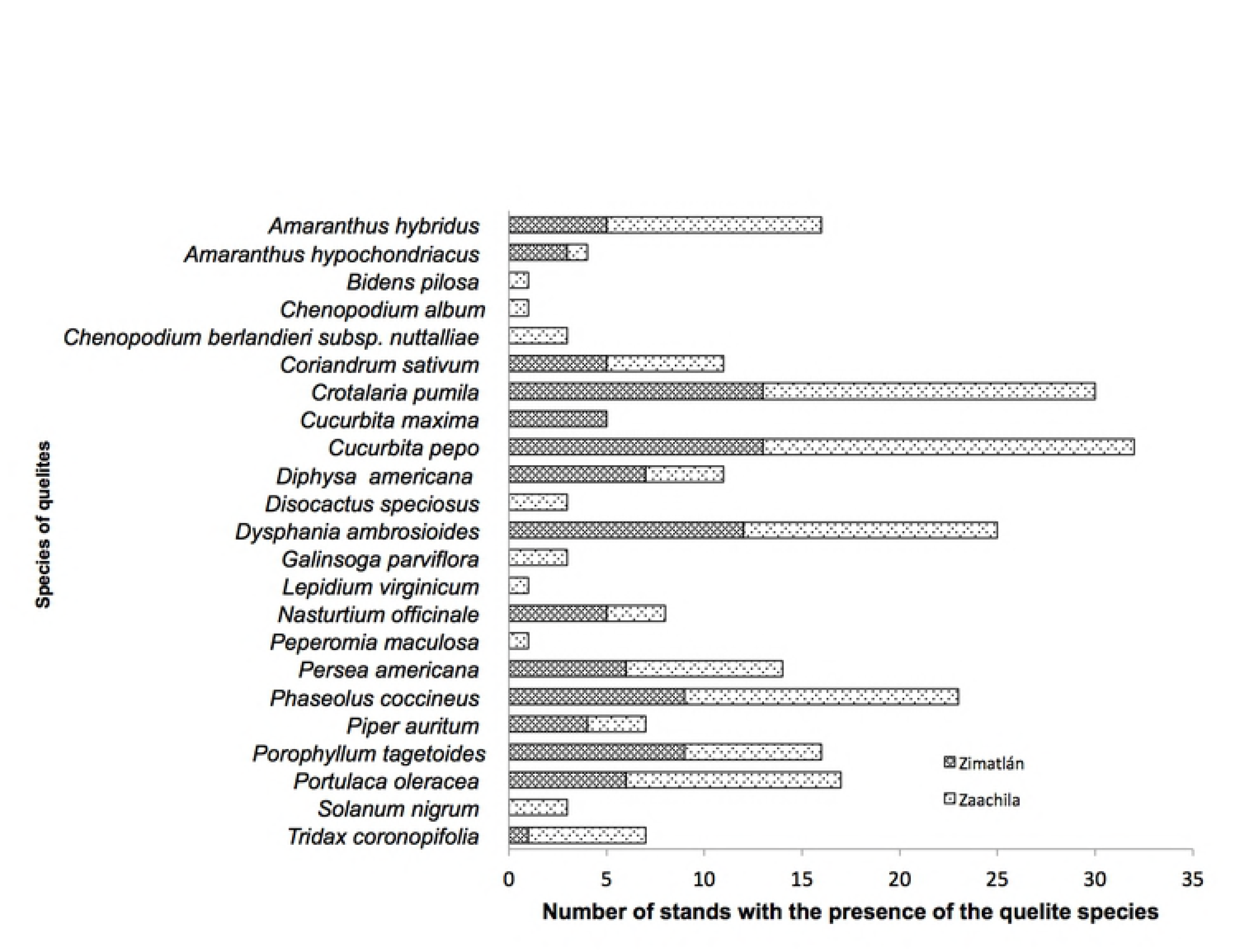
Frequency of the presence of the quelite species commercialized in the markets of Zaachila and Zimatlán.

**Table 1.** List of presence of quelite species in the markets of Zimatlán and Zaachila.

| Botanical family | Scientific name | Common name | Used part | Zi | Za |
| --- | --- | --- | --- | --- | --- |
| Amaranthaceae | *Amaranthus hybridus* L. | *quintoniles*<sup>(1)(2)</sup> | leaves | x | x |
| Amaranthaceae | *Amaranthus hypochondriacus* L. | amaranth<sup>(1)</sup>, amaranth leaf<sup>(2)</sup>,<br>*quintonil*<sup>(2)</sup> | leaves | x | x |
| Asteraceae | *Bidens pilosa* L. | *aceitillo*<sup>(2)</sup> | leaves |  | x |
| Amaranthaceae | *Chenopodium album* L. Bosc ex Moq. | *quelite cenizo*<sup>(2)</sup> | leaves |  | x |
| Amaranthaceae | *Chenopodium berlandieri* subsp.<br>*nuttalliae* (Saff.) H.D.Wilson & Heiser | *huauzontle*<sup>(2)</sup> | leaves, branches,<br>flowers, seeds |  | x |
| Apiaceae | *Coriandrum sativum* L. | coriander<sup>(1)(2)</sup> | leaves, stems | x | x |
| Fabaceae | *Crotalaria pumila* Ortega | *chepiles*<sup>(1)(2)</sup> | leaves, flowers | x | x |
| Cucurbitaceae | *Cucurbita maxima* Duchesne | pumpik flowers<sup>(1)(2)</sup>,<br>*guías de calabaza*<sup>(1)(2)</sup> | leaves, stems, flowers,<br>fruits | x | x |
| Cucurbitaceae | *Cucurbita pepo* L. | pumpik flowers<sup>(1)(2)</sup>,<br>*guías de calabaza*<sup>(1)(2)</sup> | leaves, stems, flowers,<br>fruits | x | x |
| Fabaceae | *Diphysa americana* (Mill.) M.Sousa | *guachepil*<sup>(1)(2)</sup>, *quelites de flores de*<br>*guachepil*<sup>(1)</sup>, *flor de cuachepil*<sup>(2)</sup>,<br>*Yaviza*<sup>(2)</sup> | flowers | x | x |
| Cactaceae | *Disocactus speciosus* (Cav.) Barthlott | *flor de junco*<sup>(2)</sup> | flowers |  | x |
| Amaranthaceae | *Dysphania ambrosioides* (L.)<br>Mosyakin & Clemants | *epazote*<sup>(1)(2)</sup> | leaves, stems | x | x |
| Asteraceae | *Galinsoga parviflora* Cav. | *piojito*<sup>(2)</sup>, *hierba de piojito*<sup>(2)</sup> | leaves, stems, flowers |  | x |
| Brassicaceae | *Lepidium virginicum* L. | *pierna de vieja*<sup>(2)</sup> | leaves, stems |  | x |
| Brassicaceae | *Nasturtium officinale* R.Br. | *berros*<sup>(1)(2)</sup> | leaves | x | x |
| Piperaceae | *Peperomia maculosa* (L.) Hook | *oreja de león*<sup>(2)</sup> | leaves |  | x |
| Lauraceae | *Persea americana* Mill. | leave of avocado<sup>(1)(2)</sup> | leaves | x | x |
| Fabaceae | *Phaseolus coccineus* L. | *flor de frijolón*<sup>(1)(2)</sup>, *quelite de frijolón*<sup>(1)(2)</sup>, *quelite de flor de frijolón*<sup>(2)</sup> | flowers | x | x |
| Piperaceae | *Piper auritum* Kunth. | *hoja de hierba santa*<sup>(2)</sup>, *hierba santa*<sup>(2)</sup> | leaves | x | x |
| Asteraceae | *Porophyllum tagetoides* (Kunth) DC. | *chepiches*<sup>(1)(2)</sup>, *tepiches*<sup>(1)(2)</sup> | leaves | x | x |
| Portulacaceae | *Portulaca oleracea* L. | *verdolagas*<sup>(1)(2)</sup> | leaves, stems | x | x |
| Solanaceae | *Solanum nigrum* L. | *hierba mora*<sup>(2)</sup> | leaves |  | x |
| Asteraceae | *Tridax coronopifolia* (Kunth) Hemsl. | *hierba de conejo* <sup>(1)(2)</sup> | leaves | x | x |

Markets: Zimatlán (1), Zaachila (2)

Markets: Zimatlán (1), Zaachila (2)

The most sold species belong to the botanical families *Fabaceae* and *Cucurbitaceae*, which corresponds to the species with the highest frequency in the markets (**Figure 4**). The frequency of the *quelites* in the markets of Zaachila and Zimatlán varies in the different months of the year. During most of the rainy season (June, July and August) the highest number of species was registered, *Crotalaria pumila* (*chepil*), *Cucurbita pepo* (pumpkin flower), and *Dysphania ambrosioides* (*epazote*) which were present throughout the year as shown in **Table 2**, and where are shown the species of *quelites* sold during 2017 in both markets.

**Figure 4.**
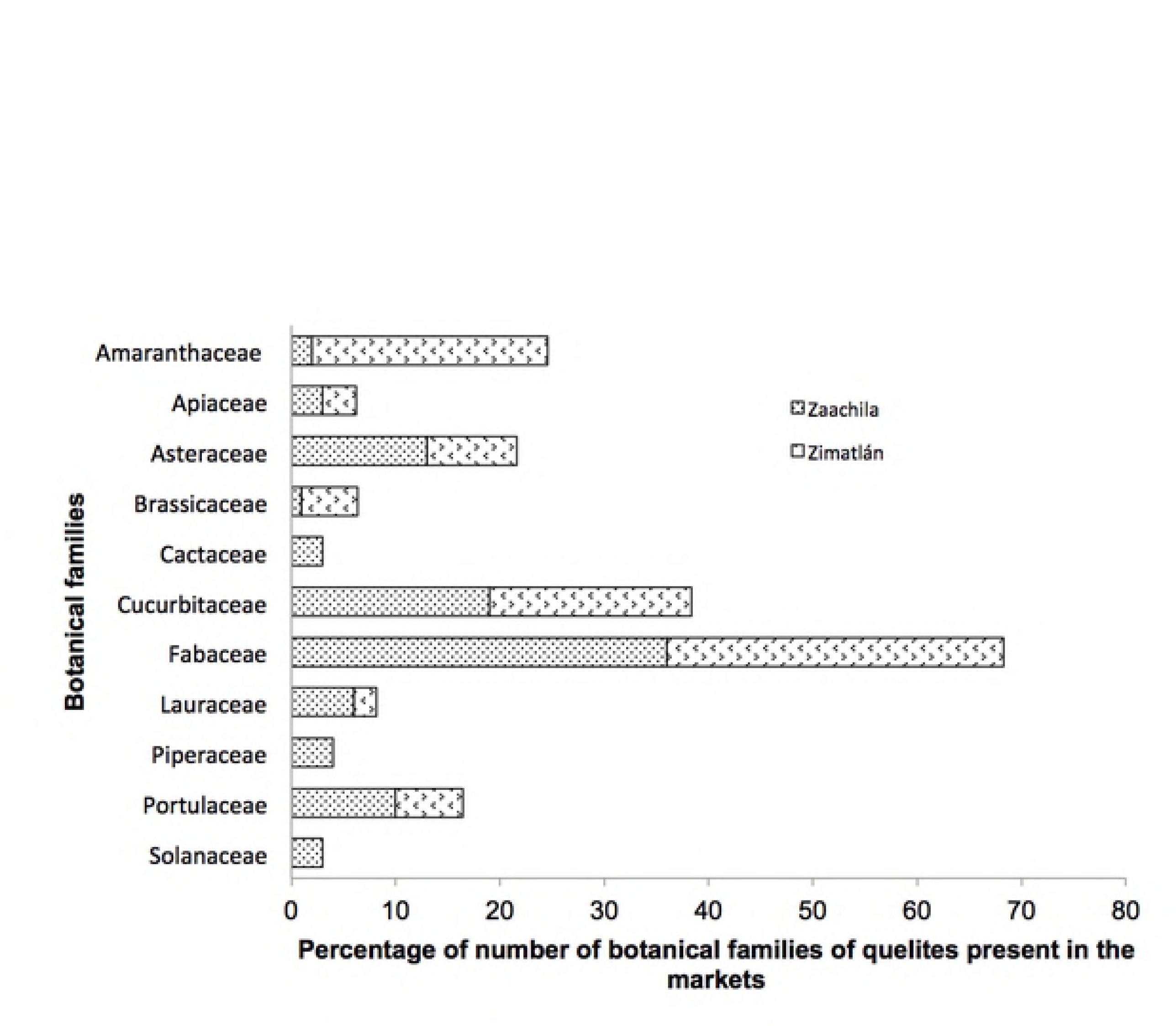
Frequency of the botanical families present in the stands of the two markets Zaachila and Zimatlán.

**Table 2.**
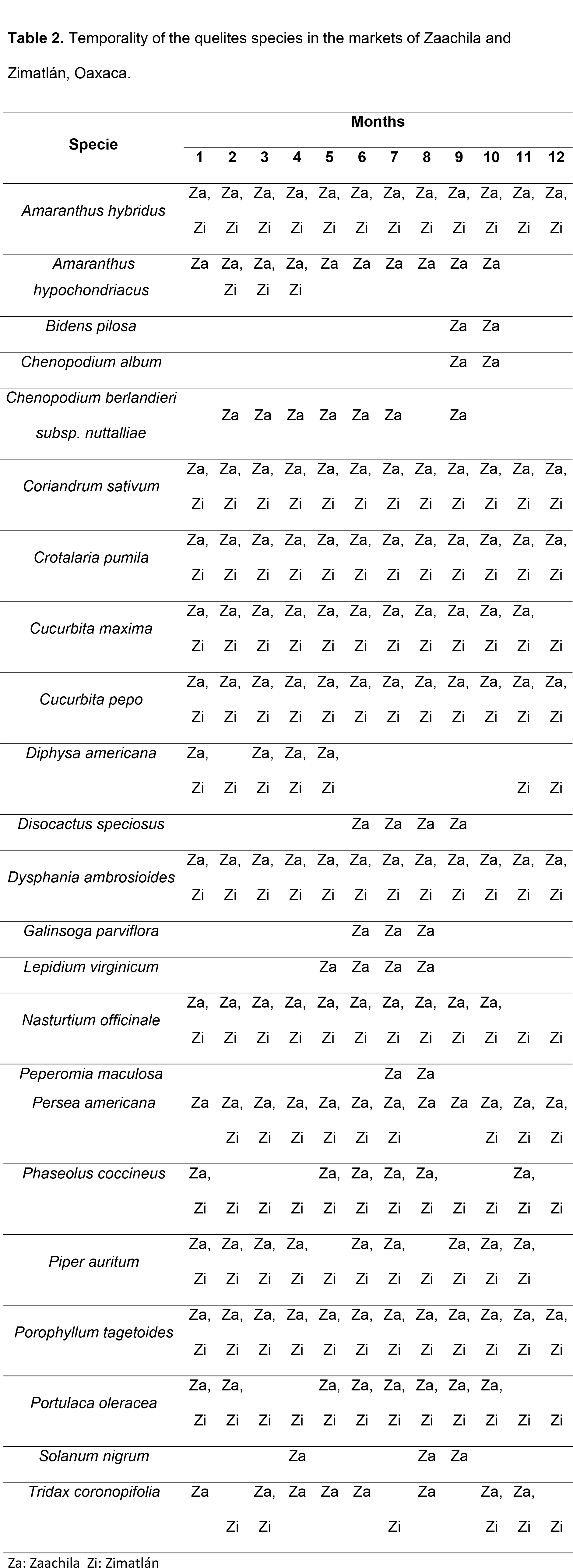
Temporality of the quelites species in the markets of Zaachila and Zimatlán, Oaxaca.

Za: Zaachila Zi: Zimatlán

## Nutritional study

In **Table 3**, are reported the results obtained from the proximal analysis performed on the floral structures of *Diphysa americana* (Q1) and *Phaseolus coccineus* (Q2).

**Table 3.** Proximal composition (%) of the *quelites* expressed in fresh matter.

| Composition | Q1 | Q2 | *p* |
| --- | --- | --- | --- |
| Moisture | 87.70 ± 0.40 | 84.55 ± 1.64 | 0.032* |
| Ash | 1.57 ± 0.06 | 1.45 ± 0.16 | 0.310 |
| Total lipids | 0.05 ± 0.00 | 0.06 ± 0.00 | 0.116 |
| Raw fiber | 1.66 ± 0.06 | 2.43 ± 0.26 | 0.008* |
| Proteins | 2.66 ± 0.10 | 3.29 ± 0.35 | 0.040* |
| Carbohydrates (NFE) | 6.36 ± 0.19 | 8.22 ± 0.87 | 0.023* |
| Energy (kJ/100g) | 152.9 ± 4.8 | 192.5 ± 20.7 | 0.027* |

Q1: *Diphyssa americana*; Q2: *Phaseolus coccineus*, NFE: nitrogen-free extract.

Q1: *Diphysa americana*; Q2: *Phaseolus coccineus,* NFE: nitrogen-free extract. Values are the mean  standard deviation of three repetitions

In relation to raw fiber, proteins, carbohydrates and energy content, the Q2 sample showed higher amounts than Q1 (p<0.05).

The results obtained for the phenolic compounds, flavonoids total and antioxidant capacity are shown in **Table 4**. It is observed that Q1 has a higher content of phenols, flavonoids and antioxidant capacity compared to Q2 (p <0.05).

**Table 4.**
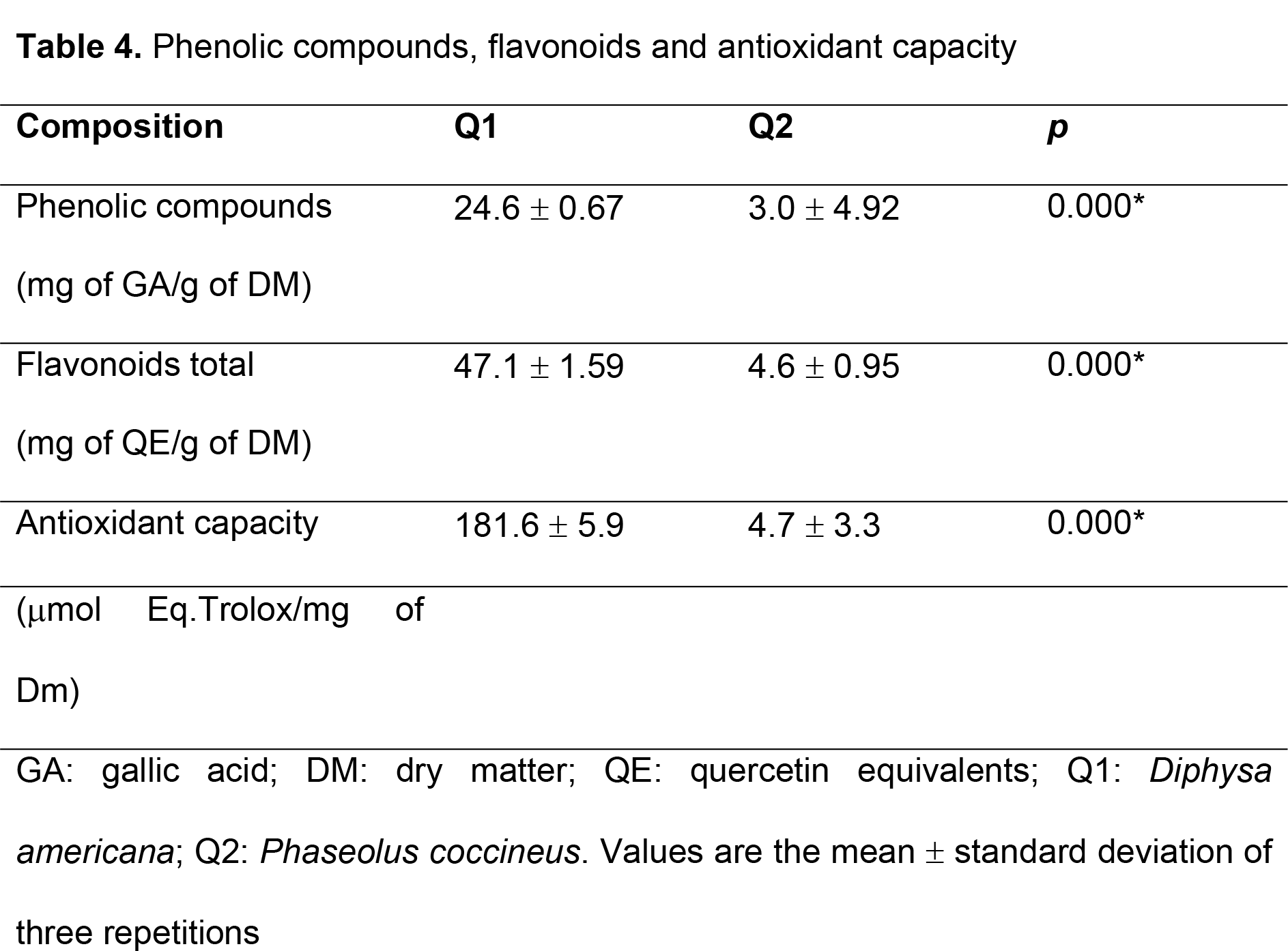
Phenolic compounds, flavonoids and antioxidant capacity.

GA: gallic acid; DM: dry matter; QE: quercetin equivalents; Q1: *Diphysa americana*; Q2: *Phaseolus coccineus*. Values are the mean ± standard deviation of three repetitions

The mineral composition of the flowers studied, reported on dry basis, is given in **Table 5**. The microelements found for *Diphysa americana* (Q1) and *Phaseolus coccineus* (Q2) were zinc, copper and Iron, while the macroelements for both samples were calcium and magnesium. As you can see, these edible flowers are an excellent source of minerals, especially calcium and magnesium. The content of these macroelements are in the range of 344.88 to 1433.91 mg/100 g and 47.23 to 146.7 mg/100 g, respectively. Being significantly higher (p<0.05) for Q2 than for Q1. While zinc and copper occur in greater quantities in Q1 than in Q2.

**Table 5.** Mineral composition (mg/100 g) of the plants on a dry basis.

| Minerals | Q1 | Q2 | *p* |
| --- | --- | --- | --- |
| Zinc | 4.25 ± 0.24 | 5.00 ± 0.13 | 0.009* |
| Copper | 0.99 ± 0.05 | 0.52 ± 0.03 | 0.000* |
| Iron | 10.98 ± 0.48 | 11.71 ± 0.6 | 0.196 |
| Calcium | 344.88 ± 7.19 | 1433.91 ± 30.22 | 0.000* |
| Magnesium | 47.23 ± 1.77 | 146.70 ± 14.70 | 0.000* |

Q1: *Diphyssa americana*; Q2: *Phaseolus coccineus*. Values are the mean ± standard deviation of three repetitions

Q1: *Diphysa americana*; Q2: *Phaseolus coccineus*. Values are the mean ± standard deviation of three repetitions

## DISCUSSION

### Frequency of species considered *quelites* in the markets of Villa de Zaachila and Zimatlán de Álvarez

Eight species of *quelites* such as *aceitillo* (*Bidens pilosa*), *quelite cenizo* (*Chenopodium album*), *huauzontle* (*Chenopodium belandieri* ssp *nuttalie*), *flor de junco* (*Disocactus speciosus*), *hierba de piojito* (*Galinsoga parviflora*), *pierna de vieja* (*Lepidium virginicum*), *oreja de león* (*Peperomia maculosa*) and *hierba mora* (*Solanum nigrum*), are only sold in the market of Zaachila. This richness and diversity of species coincide with the information documented by Cortés Romero (2005) [37]; López-Aguilar (2005) [38]; Tapia *et al*. (2012) [39]; Martínez-Bolaños (2014) [40], who point out that in the stands outside this market there is a greater influx of sellers than in other markets in the Valles Centrales’ region. The people who sell the *quelites* in these markets come from different places in the Valles Centrales’ region, which is divided into seven Districts, thus showing an important cultural diversity. In the market of Zimatlán the *quelite* sellers come from three Districts of the Central Valleys: 1) from the District of Zimatlán come sellers from the municipalities of Zimatlán de Álvarez, San Bernardo Mixtepec, San Miguel Mixtepec and Ayoquezco de Aldama; 2) of the District of Ocotlán come sellers from the municipalities of Santa Lucia Ocotlan, San Antonino Castillo Velasco, San Sebastian Teitipac and Ocotlan de Morelos; and 3) of the District of Tlacolula come only from the municipality of Tlacolula de Matamoros. In the Zaachila market, sellers come from four Districts: 1) from the Central District, mainly sellers from the municipality of Cuilapam de Guerrero and from a town in the same municipality called El Carrizal; 2) of the District of Ocotlán arrive from the municipalities of San Antonino Castillo Velasco; 3) of the District of Zimatlán arrive from the municipality of Zimatlán de Álvarez; and 4) of the District of Zaachila come from the municipalities of Santa Inés del Monte and Villa de Zaachila, as shown in **Figure 1**. Occasionally people arrive from the region of the Coast, as was the case of a person from the municipality Candelaria Loxicha belonging to the District of Pochutla. Observing people from different places, including a coastal municipality, supports the cultural relevance of this market in the sale of *quelites* and native products of the region.

*Cucurbita pepo* is the plant with the highest frequency in these two markets, followed by *Crotalaria pumila* and *Dysphania ambrosioides*, probably because of the first species are also sold the stems, flowers and fruits, to cook the famous “*sopa de guías*”, a very emblematic dish in Oaxacan cuisine; In addition, the pumpkin is grown in orchards and fields throughout the year. The *quelites* that are sold in these markets are eaten both fresh and dried and the main edible parts are the leaves, except for the species selected for the chemical-nutrimental study, where the botanical structure that is eaten are the flowers.

### Nutritional study

The moisture content of the *quelites* studied is similar to those reported by Mohammad et al. 2007 [41], which were found for *Amaranthus viridis* L., *Chenopodium murale* L., *Nasturtium officinale R.Br*., and *Scandix pecten-veneris* L., a moisture range of 81.3% to 90.5%. The values of the ash is similar to that reported for *Nasturtium officinale*, (1.77%). The protein content of the analyzed samples was high for the two *quelites* analyzed (2.66% for Q1 and 3.29% for Q2). The high content of raw protein in these plants can be of great importance to be incorporated in the diets of marginalized communities with problems of malnutrition. In addition, the fat content is very low (<0.1%) in both samples, even compared to that reported in other wild and edible plants [41], where they report approximately 0.5%. The raw fiber content was 1.66% for Q1 and 2.43% for Q2. The World Health Organization establishes the consumption of 25 to 30 g of fiber per day, which, although it does not contribute to the nutritional value of food, helps maintain a good digestion of the organism. Therefore, the high levels of fiber contained in flowers make them attractive to integrate into the diet of people. The Q2 *quelite* sample showed the highest levels of carbohydrates (8.22%) and therefore an energy content of 192.5 kJ/100 g of dry sample. According to the Food Guide for the Mexican population, a serving of vegetables equals 104.6 kcal, 4 g of carbohydrates, 0 g of fat and 2 g of protein, so the equivalence of the *quelites* would be that 50 g of *quelites* to provide between 77.4 to 96.2 kcal, 3.2 to 4.1 g of carbohydrates, 0 g of fats and 1.33 to 1.7 g of proteins.

By showing a relationship between phenolic compounds and the elimination of free radicals, the phenolic content of plants greatly contributes to their antioxidant potential [42]. In 12 species, different from those studied in this work, evaluated edible flowers found a concentration of phenolic compounds in a range of 2.53 to 5.28 mg of gallic acid/g of fresh matter (26.4 to 45.7 mg of gallic acid/g of dry matter) [15]. In this sense, the *Diphysa americana* (Q1) *quelite* species is around that reported in the literature. DPPH is a rapid and sensitive method to evaluate the antioxidant capacity of plant extracts. This is based on the discoloration of DPPH in the presence of antioxidants in the samples analyzed. According to Shahidi and Wanasundara, 1992 [43] the antioxidant power is mainly due to the redox nature of the phenolic compounds. Our results also confirm the aforementioned. The results obtained from the total content of phenols and flavonoids total for the Q1 sample also show higher activity to trap free radicals.

The content of the macroelements, calcium and magnesium, are in the range of 344.88 to 1433.91 mg/100 g and 47.23 to 146.7 mg/100 g, respectively. The literature reports on 12 species of flowers a range of 23.9 to 49.2 mg/100 g of fresh matter (FM) for calcium (285.6 to 425.9 mg/100 g of dry matter) and from 10.5 to 20.5 mg/100 g of FM for magnesium (110.0 to 212.0 mg/100 g of dry matter) [15]. This indicates that Q1 and Q2 contain more calcium but less magnesium tan previously reported for 12 flowers in the literature. In the case of microelements, the quantification of zinc in Q1 and Q2 (4.25 mg/100 g and 5.00 mg/100 g of dry matter, respectively) is higher than that reported for species of *quelites* such as *Nasturtium officinale* but below that for *Amaranthus viridis* L. [41]. Zinc is a trace mineral that is especially important for the normal functioning of the immune system. In edible flower species such as *Rosa odorata*, *Tropaeolum majus* and *Tagetes patula*, has been reported a concentration of 4.5 to 13.7 mg/100 g zinc of dry matter. For flowers of the species *Begonia boliviensis*, *Chrysanthemum frutescens* and *Dianthus caryophyllus* the reported iron content is in the range of 1.87 to 8.53 mg/100 g of dry matter, while for *Centaurea cyanus*, *Antirrhinum majus* and *Dianthus caryophyllus*, the content of copper has been in the range of 0.91 to 2.49 mg/100 g of dry matter [15]. In **Table 4** it can be observed that the two species studied *Diphysa americana* (Q1) and *Phaseolus coccineus* (Q2) are well above of what was previously reported in iron. Metal ions are crucial for our body, they frequently serve as cofactors in enzymatic reactions. They are also important for the maintenance of the protein structure.

One third of the body’s proteins bind to metal ions, requiring approximately 10% of the enzymes in our body zinc for its activity [44]. Zinc deficiency contributes to the death of 800,000 children worldwide per year [45]. In general, the deficiency of copper, zinc and manganese damages the cellular function, affects the development and growth, the immune system and metabolism. Copper is the main constituent of cuproproteins which are involved in energy production, iodine metabolism and metabolism of neurotransmitter synthesis [46]. Iron deficiency induces major disorders such as anemia, which affects about a third of the world’s population [47]. The recommended dietary intake (RDA) in the USA is 11 mg/day for men and 8 mg/day for zinc women; 900 ug/day of copper for both sexes and 8 mg/day for men and 18 mg/day for women of iron. So when the *quelites Diphysa americana* and *Phaseolus coccineus* are properly incorporated into the human diet, the daily requirements would be covered with native plants of the region.

## CONCLUSIONS

The use of a large number of wild species considered *quelites* in the markets of the Valles Centrales’ region of Oaxaca has a pre-Hispanic origin and its consumption still continues on a daily basis in rural and urban communities, where there is a wide diversity of wild resources for local gastronomy and other ways of use. The sale of quelite species is a very important activity that complements the incomes of local farmers, mainly of peasant women and knowledgeable about the management of agroecosystems such as family gardens, milpa and sometimes the fields. You can clearly tell the difference between the retail and wholesale, the latter being the one that is suggested to be more economically profitable. Both markets sell different species in the wild and grown state throughout the year. Regarding the chemical study of the two species of *quelites* have shown to be a rich source of proteins and bioactive compounds. In particular, *Diphysa americana* was the one that showed the highest content of phenolic compounds, flavonoids and, therefore, greater antioxidant capacity, suggesting an alternative for future research in relation to the different forms of preparation and incorporation into the feeding patterns of the population. This is the first report on the chemical study about of *Diphysa americana* and *Phaseolus coccineus* flowers.

## Bibliography

1. Toledo MV. La diversidad biológica de México. Ciencia y Desarrollo. 1988;14(81):17-30.

2. Rzedowski J. Vegetación de México. 1ra. Edición digital, Comisión Nacional para el Conocimiento y Uso de la Biodiversidad, México. 2006:504.

3. Miranda F, Hernández-X E. Los tipos de vegetación en México y su clasificación. Edición Conmemorativa (1963-2013) ed: Sociedad Botánica de México. Comisión Nacional para el Conocimiento de la Biodiversidad. Fondo de Cultura Económica; 2014.

4. García-Mendoza AJ, Meave JA. Diversidad florística de Oaxaca: de musgos a angiospermas (colecciones y lista de especies). Universidad Nacional Autónoma de México-Comisión Nacional para el Conocimiento y Uso de la Biodiversidad. México. 2011:352.

5. Linares E, Aguirre J. Los quelites, un tesoro culinario. Universidad Nacional Autónoma de México. Instituto Nacional de la Nutrición “Salvador Zubirán”. Instituto de Biología. 1992:143.

6. Parrott N, Wilson N, Murdoch J. Spatializing Quality: Regional Protection and the Alternative Geography of Food. European Urban and Regional Studies. 2002;9(3):241-61.

7. González-Mendoza D, Ascencio-Martínez D, Hau Poox A, Méndez-Trujillo V, Grimaldo-Juarez O, Santiaguillo-Hernández JF, et al. Phenolic compounds and physicochemical analysis of Physalis ixocarpa genotypes. Scientific Research and Essays. 2011;6(17):3808-14.

8. Bergier K, Kuzniak E, Sklodowska M. Antioxidant potential of Agrobacterium transformed and non transformed Physalis ixocarpa plants grown in vitro and ex vitro. Postepy Hig Med Dosw (Online). 2012;66:976-82.

9. Casas A, Viveros JL, Caballero J. Etnobotánica mixteca: sociedad, cultura y recursos naturales en la Montaña de Guerrero, México, In: Artes. Instituto Nacional Indigenista. Consejo Nacional para la Cultura y las Artes. 1994:336.

10. Caballero J, Casas A, Cortés L, Mapes C. Patrones en el conocimiento, uso y manejo de plantas en pueblos de México. E. Atacameños, Editor. 1998;16:1-15.

11. Caballero J, Rendón B, Rebollar S, Martínez MA. Uso y Manejo tradicional de los Recursos Vegetales en México. In: Estudio sobre la relación entre seres humanos y plantas en los albores del sigloo XXI. 2001:79-100.

12. Linares E, Bye R. “Naturaleza e identidad nacional”, un Elogio de la Cocina Mexicana, Patrimonio Cultural de la Humanidad, México: Conservatorio de la Cultura Gastronómica Mexicana S.C. y Artes de México. 2012:57-67.

13. Flores EM, Marín WA. Diphysa americana (Mill.) M. Sousa. In: Tropical Tree Seed Manual. (J. A. Vozzo, Editor). https://rngr.net/publications/ttsm. United States Department of Agriculture Forest Service. 2001:439-41 pp.

14. Ruiz Salazar RAdldgdPcLdlsdcHdMTdMeCeB. Análisis de la diversidad genética de Phaseolus cocineus L. de la subprovincia de cuarso Husteco de México. Tesis de Maestría en Ciencias en Biotecnología Genómica. Centro de Biotecnología Genómica, Instituto Politécnico Nacional, Reynosa, Tamaulipas. 2009:65.

15. Rop O, Mlcek J, Jurikova T, Neugebauerova J, Vabkova J. Edible flowers‐‐a new promising source of mineral elements in human nutrition. Molecules. 2012;17(6):6672-83.

16. Kopec K, Balik J. Kvalitologie Zahradnickych Produktu, 1st ed.; Mendel University of Agriculture and Forestry in Brno: Brno, Czech Republic 2008.

17. Yang SL, Walters TW. Ethnobotany and the economic role of the Cucurbitaceae in China. Econ Bot 1992;46:349-67.

18. Upadhyay RK. Kareel plant: A natural source of medicines and nutrients. Int J Green Pharm. 2011;5:255-65.

19. Aletor O, Oshodi AA, Ipinmoroti K. Chemical composition of common leafy vegetables and functional properties of their leaf protein concentrates. Food Chem 2002;78:63-8.

20. Lin CC, Chung YC, Hsu CP. Potential roles of longan flower and seed extracts for anti-cancer. World J Exp Med. 2012;2(4):78-85.

21. Hsu CL, Fang SC, Yen GC. Anti-inflammatory effects of phenolic compounds isolated from the flowers of Nymphaea mexicana Zucc. Food Funct. 2013;4(8):1216-22.

22. Besbes-Hlila M, Omri A, Ben-Jannet H, Lamari A, Aouni M, Selmi B. Phenolic composition, antioxidant and anti-acetylcholinesterase activities of the tunisian scabiosa arenaria. Pharm Biol 2013;51(5):525-32.

23. Friedman H, Agami O, Vinokur Y, Droby S, Cohen L, Refaeli G, et al. Characterization of yield, sensitivity to Botrytis cinerea and antioxidant content of several rose species suitable for edible flowers. Sci Hortic 2010;123:395-401.

24. Mato M, Onazaki T, Ozeki Y, Higeta D, Itoh Y, Yoshimoto Y, et al. Flavonoid biosynthesis in white flowered sim carnations (Dianthus caryophyllus). Sci Hortic 2000;84:333-47.

25. INEGI. Prontuario de información geográfica municipal de los Estados Unidos Mexicanos. Villa de Zaachila, Oaxaca. Clave geoestadística 20565. 2005.

26. INEGI. Prontuario de información geodráfica municipal de los Estados Unidos Mexicanos. Zimatlán de Álvarez, Oaxaca. Clave geoestadística 20570. 2005.

27. Lot A, Chiang F. Manual de herbario. Administración y manejo de colecciones, técnicas, recolección y preparación de ejemplares botánicos. Consejo Nacional de la Flora de México, A.C. IBUNAM. 1990:142.

28. Manzanero-Medina GI, Flores-Martínez E, Sandoval Z, Bye R. Etnobotánica de siete raíces medicinales en el Mercado de Sonora de la Ciudad de México. Polibotánica. 2009(27):191-228.

29. AOAC OMoA. Gaithersburg, Maryland, USA, AOAC International. 17 th ed 2000.

30. FAO. Food and Nutrition paper. Food energy-methods of analysis and conversion factors. Rome December 2001.

31. Rababah TM, Hettiarachchy NS, Horax R. Total phenolics and antioxidant activities of fenugreek, green tea, black tea, grape seed, ginger, rosemary, gotu kola, and ginkgo extracts, vitamin E, and tert-butylhydroquinone. J Agric Food Chem. 2004;52(16):5183-6.

32. Singleton VL, Rossi JA. Colorimetry of total phenolics with phosphomolybdic–phosphotungstic acid reagents. American Journal of Enology and Viticulture. 1965;16:144-58.

33. Chen L, Xin X, Yuan Q, Su D, Liu W. Phytochemical properties and antioxidant capacities of various colored berries. Journal of the Science of Food and Agriculture. 2014;94(2):180-8.

34. Matthaus B. Antioxidant activity of extracts obtained from residues of different oilseeds. J Agric Food Chem. 2002;50(12):3444-52.

35. Dürust N, Sumengen D. Ascorbic acid and element contents of foods of Trabzon (Turkey). Journal of Agricultural Food Chemistry. 1997;45(6):2085-7.

36. Julián-Loaeza AP, Santos-Sánchez NF, Valadez-Blanco R, Sánchez-Guzmán BS, Salas-Coronado R. Chemical composition, color, and antioxidant activity of three varieties of Annona diversifolia Safford fruits. Industrial Crops and Products. 2011;34:1262-8.

37. Cortés-Romero O. Análisis comparativo de la diversidad de especies suculentas en mercados de los Valles Centrales de Oaxaca. Memoria de Residencia Profesional, México: Instituto Tecnológico del Valle de Oaxaca. 2005:128.

38. López-Aguilar P.. Diversidad y uso de las plantas silvestres en los mercados del Valle de Oaxaca. Memoria de Residencia Profesional Licenciatura en Biología. Instituto Tecnológico Agropecuario de Oaxaca. 2005:66.

39. Tapia PD, Manzanero-Medina GI, Flores MA. Etnobotánica de plantas silvestres en mercados tradicionales. Región de Valles Centrales, Oaxaca, México. Editorial Académica Española. 2012:73.

40. Martínez-Bolaños KA. El valor de uso de las plantas ornamentales-rituales comercializadas en los mercados de los Valles Centrales del estado de Oaxaca. Maestría en Ciencias en Conservación y Aprovechamiento de Recursos Naturales. CIIDIR-IPN-Oaxaca. 2014:49.

41. Mohammad I, Farah NT, Muhammad IJ, Asifullah K, Ikhtiar K. Analysis of Nutritional Components of Some Wild Edible Plants. Jour Chem Soc Pak. 2007.

42. Zovko Koncic M, Kremer D, Karlovic K, Kosalec I. Evaluation of antioxidant activities and phenolic content of Berberis vulgaris L. and Berberis croatica Horvat. Food Chem Toxicol. 2010;48(8-9):2176-80.

43. Shahidi F, Wanasundara PK. Phenolic antioxidants. Crit Rev Food Sci Nutr. 1992;32(1):67-103.

44. Azia A, Levy R, Unger R, Edelman M, Sobolev V. Genome-wide computational determination of the human metalloproteome. Proteins. 2015;83(5):931-9.

45. Hagan LL, Nti C, G.M M, Danquah A. Men’s Involvement in Caregivers’ Participation in ENAM Project Micro-credit Programme and Children’s Animal Source Food Consumption in Rural Ghana. Research Brief 10-04-ENAM. Global Livestock Collaborative Research Support Program (GL-CRSP). University of California-Davis, Davis, CA. 2010.

46. Mejia-Rodriguez F, Shamah-Levy T, Villalpando S, Garcia-Guerra A, Mendez-Gomez Humaran I. Iron, zinc, copper and magnesium deficiencies in Mexican adults from the National Health and Nutrition Survey 2006. Salud Publica Mex. 2013;55(3):275-84.

47. Senesse P, Meance S, Cottet V, Faivre J, Boutron-Ruault MC. High dietary iron and copper and risk of colorectal cancer: a case-control study in Burgundy, France. Nutr Cancer. 2004;49(1):66-71.

